# ERGA-BGE genome of *Stigmatoteuthis arcturi* Robson, 1948: the jewelled squid

**DOI:** 10.1101/2025.01.13.632676

**Authors:** Fernando Ángel Fernández-Álvarez, Ainhoa Bernal-Bajo, Nuria Escudero, María Conejero, Ana Riesgo, Rosa Fernández, Rita Monteiro, Astrid Böhne, Laura Aguilera, Marta Gut, Tyler S. Alioto, Francisco Câmara Ferreira, Fernando Cruz, Jèssica Gómez-Garrido, Tom Brown

## Abstract

The *Stigmatoteuthis arcturi* reference genome offers a valuable resource for understanding the evolutionary patterns of oceanic squids, who perform important ecological roles as both predators and prey in mesopelagic and deep-sea environments, while their genomes remain understudied. The entirety of the genome sequence was assembled into 46 contiguous chromosomal pseudomolecules. This chromosome-level assembly encompasses 3.25 Gb, composed of 1,268 contigs and 497 scaffolds, with contig and scaffold N50 values of 11.0 Mb and 74.6 Mb, respectively.

## Introduction

*Stigmatoteuthis arcturi* Robson, 1948 belongs to the family Histioteuthidae Verrill, 1880–1881, known as the jewelled squids, which are one of the most important components of the diet of endangered megafauna, such as sperm whales (Clarke, 2006). Jewelled squids are characterised by a unique morphology with a number of photophores on their skin to break its shadow and deceive predators from deeper waters. They also possess high levels of asymmetry in their bodies, peaking in a large difference in size, morphology and pigmentation of their eyes, themselves specialised to different tasks (Thomas et al., 2017). While the larger left eye looks at the dim light coming from the surface to spot their megafaunal predators, the smaller right eye looks towards the bottom looking for bioluminescence of their micronekton prey. *S. arcturi* is one of the three allopatric species of the genus *Stigmatoteuthis* Pfeffer, 1900, which are characterised by duplicated terminal parts of the male reproductive system and with subtle morphological differences among them, which can only be recognised in mature males (Young & Vecchione, 2016). It distributes in tropical and subtropical Atlantic offshore mesopelagic waters and as with any other cephalopod, *S. arcturi* is fast growing, fueled by a very intense predatory activity. Jewelled squids are paratenic hosts of parasitic ascarid helminths, such as *Anysakis* Dujardin, 1845 and other nematodes (Palomba et al., 2021). They transfer these parasites to higher trophic level hosts, such as commercially important swordfishes and endangered toothed whales, where these parasites finish their life cycles.

Jewelled squids are the largest component of sperm whales diets in many areas (Clarke, 2006) and an important prey item for many other megafaunal species, such as dolphins, sharks and swordfishes. Thus, they represent an important energy and biomass link among lower and upper trophic levels in pelagic oceanic food webs. Jewelled squids have an unsettled position in the phylogeny of oceanic squids (Fernández-Álvarez et al., 2022). The genome of *S. arcturi* is a valuable tool to solve the phylogenetic position of jewelled squids within oceanic squids.

The generation of this reference resource was coordinated by the European Reference Genome Atlas (ERGA) initiative’s Biodiversity Genomics Europe (BGE) project, supporting ERGA’s aims of promoting transnational cooperation to promote advances in the application of genomics technologies to protect and restore biodiversity (Mazzoni et al., 2023).

## Materials & Methods

ERGA’s sequencing strategy includes Oxford Nanopore Technology (ONT) and/or Pacific Biosciences (PacBio) for long-read sequencing, along with Hi-C sequencing for chromosomal architecture, Illumina Paired-End (PE) for polishing (i.e. recommended for ONT-only assemblies), and RNA sequencing for transcriptomic profiling, to facilitate genome assembly and annotation.

### Sample and Sampling Information

Ainhoa Bernal-Bajo sampled one specimen of *Stigmatoteuthis arcturi*, which was identified based on morphology by Fernando Ángel Fernández-Álvarez, from international Atlantic waters near Azores and the Canary Islands on 28th February 2023. As sampling was performed in international waters, no permit is required. Sampling was performed using a Mesopelagos net.

The specimen was euthanized by freezing at -80 ºC. Until DNA and RNA extractions, samples were preserved at -80 ºC.

### Vouchering information

Physical reference material for the here sequenced specimen has been deposited in the National Museum of Natural History of Madrid (MNCN-CSIC) https://www.mncn.csic.es/es/colecciones under the accession number MNCN 15.06/521.

Frozen reference tissue material of skin is available from the same individual at the Biobank National Museum of Natural History of Madrid (MNCN-CSIC) https://www.mncn.csic.es/es/colecciones under the ID MNCN-ADN-151723.

An electronic voucher image of the sequenced individual is available from ERGA’s EBI BioImageArchive dataset https://www.ebi.ac.uk/biostudies/bioimages/studies/S-BIAD1012?query=ERGA under accession IDs SAMEA114541340_1.jpg, SAMEA114541340_2.jpg, and SAMEA114541340_3.jpg.

### Data Availability

*S. arcturi* and the related genomic study were assigned to Tree of Life ID (ToLID) ‘xcStiArct1’ and all sample, sequence, and assembly information are available under the umbrella BioProject PRJEB81320. The sample information is available at the following BioSample accessions: SAMEA114541340, SAMEA114541342, SAMEA114541343, and SAMEA114541354. The genome assembly is accessible from ENA under accession number GCA_964276865.1. Sequencing data produced as part of this project are available from ENA at the following accessions: ERX13202607, ERX13202608, ERX13202609 and ERX13202610. Documentation related to the genome assembly and curation can be found in the ERGA Assembly Report (EAR) document available at https://github.com/ERGA-consortium/EARs/tree/main/Assembly_Reports/Stigmatoteuthis_arcturi/xcStiArct1. Further details and data about the project are hosted on the ERGA portal at https://portal.erga-biodiversity.eu/data_portal/2053936.

### Genetic Information

The estimated genome size, based on ancestral taxa, is 3.2 Gb. This is a diploid genome with a haploid number of 46 chromosomes (2n=92). All information for this species was retrieved from Genomes on a Tree (Challis et al., 2023).

### DNA/RNA processing

DNA was extracted from arm tissue using the Blood & Cell Culture DNA Midi Kit (Qiagen) following the manufacturer’s instructions. DNA quantification was performed using a Qubit dsDNA BR Assay Kit (Thermo Fisher Scientific), and DNA integrity was assessed using a Genomic DNA 165 Kb Kit (Agilent) on the Femto Pulse system (Agilent). The DNA was stored at +4ºC until used.

RNA was extracted using an RNeasy Mini Kit (Qiagen) according to the manufacturer’s instructions. RNA was extracted from four different specimen parts: muscle, skin, arm tissue and tentacle stalk. RNA quantification was performed using the Qubit RNA BR kit and RNA integrity was assessed using a Bioanalyzer 2100 system (Agilent) RNA 6000 Nano Kit (Agilent). RNA was equimolarly pooled for the library preparation and stored at -80ºC until used.

### Library Preparation and Sequencing

For long-read whole genome sequencing, a library was prepared using the SQK-LSK114 Kit (Oxford Nanopore Technologies, ONT) and was sequenced on a PromethION 24 A Series instrument (ONT). A short-read whole genome sequencing library was prepared using the KAPA Hyper Prep Kit (Roche). A Hi-C library was prepared from arm tissue using the Dovetail Omni-C kit (Cantata Bio), followed by the KAPA Hyper Prep kit for Illumina sequencing (Roche). The RNA library from the pooled sample was prepared using the KAPA mRNA Hyper prep kit for Illumina sequencing (Roche). The short-read sequencing libraries were processed on a NovaSeq 6000 instrument (Illumina). In total, 180.5 Gb Oxford Nanopore, 203.5 Gb Illumina WGS shotgun, and 195.3 Gb HiC data were sequenced to generate the assembly.

### Genome Assembly Methods

The genome was assembled using the CNAG CLAWS pipeline (Gomez-Garrido, 2024). Briefly, reads were preprocessed for quality and length using Trim Galore v0.6.7 and Filtlong v0.2.1, and initial contigs were assembled using NextDenovo v2.5.0, followed by polishing of the assembled contigs using HyPo v1.0.3, removal of retained haplotigs using purge-dups v1.2.6 and scaffolding with YaHS v1.2a. Finally, assembled scaffolds were curated via manual inspection using Pretext v0.2.5 with the Rapid Curation Toolkit (https://gitlab.com/wtsi-grit/rapid-curation) to remove any false joins and incorporate any sequences not automatically scaffolded into their respective locations in the chromosomal pseudomolecules (or super-scaffolds). Summary analysis of the released assembly was performed using the ERGA-BGE Genome Report ASM Galaxy workflow (https://doi.org/10.48546/workflowhub.workflow.1103.2).

## Results

### Genome Assembly

The genome assembly has a total length of 3,249,387,216 bp in 497 scaffolds (Figures 1 & 2), with a GC content of 34.92%. The assembly has a contig N50 of 10,980,000 bp and L50 of 88 and a scaffold N50 of 74,550,748 bp and L50 of 19. The assembly has a total of 771 gaps, totaling 154.2 kb in cumulative size. The single-copy gene content analysis using the Metazoa database with BUSCO (Manni et al., 2021) resulted in 94.8% completeness (94.0% single and 0.8% duplicated). 80.5% of reads k-mers were present in the assembly and the assembly has a base accuracy Quality Value (QV) of 35.4 as calculated by Merqury (Rhie et al., 2020).

**Figure 1.**
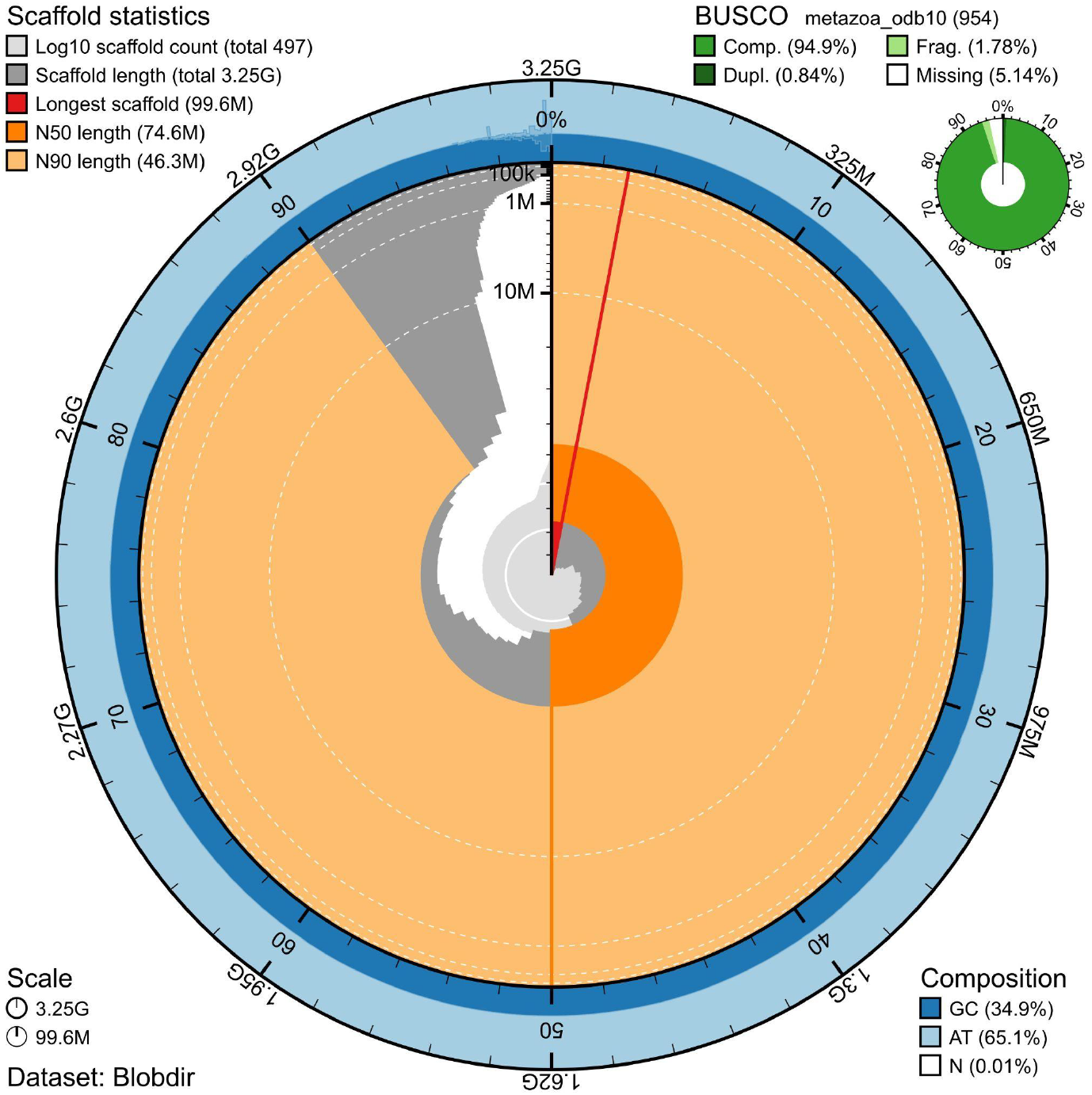
Snail plot summary of assembly statistics. The main plot is divided into 1,000 size-ordered bins around the circumference, with each bin representing 0.1% of the 3,249,387,216 bp assembly including the mitochondrial genome. The distribution of sequence lengths is shown in dark grey, with the plot radius scaled to the longest sequence present in the assembly (99,556,817 bp, shown in red). Orange and pale-orange arcs show the scaffold N50 and N90 sequence lengths (74,550,748 and 46,349,533 bp), respectively. The pale grey spiral shows the cumulative sequence count on a log-scale, with white scale lines showing successive orders of magnitude. The blue and pale-blue area around the outside of the plot shows the distribution of GC, AT, and N percentages in the same bins as the inner plot. A summary of complete, fragmented, duplicated, and missing BUSCO genes found in the assembled genome from the Metazoa database (odb10) is shown in the top right.

**Figure 2.**
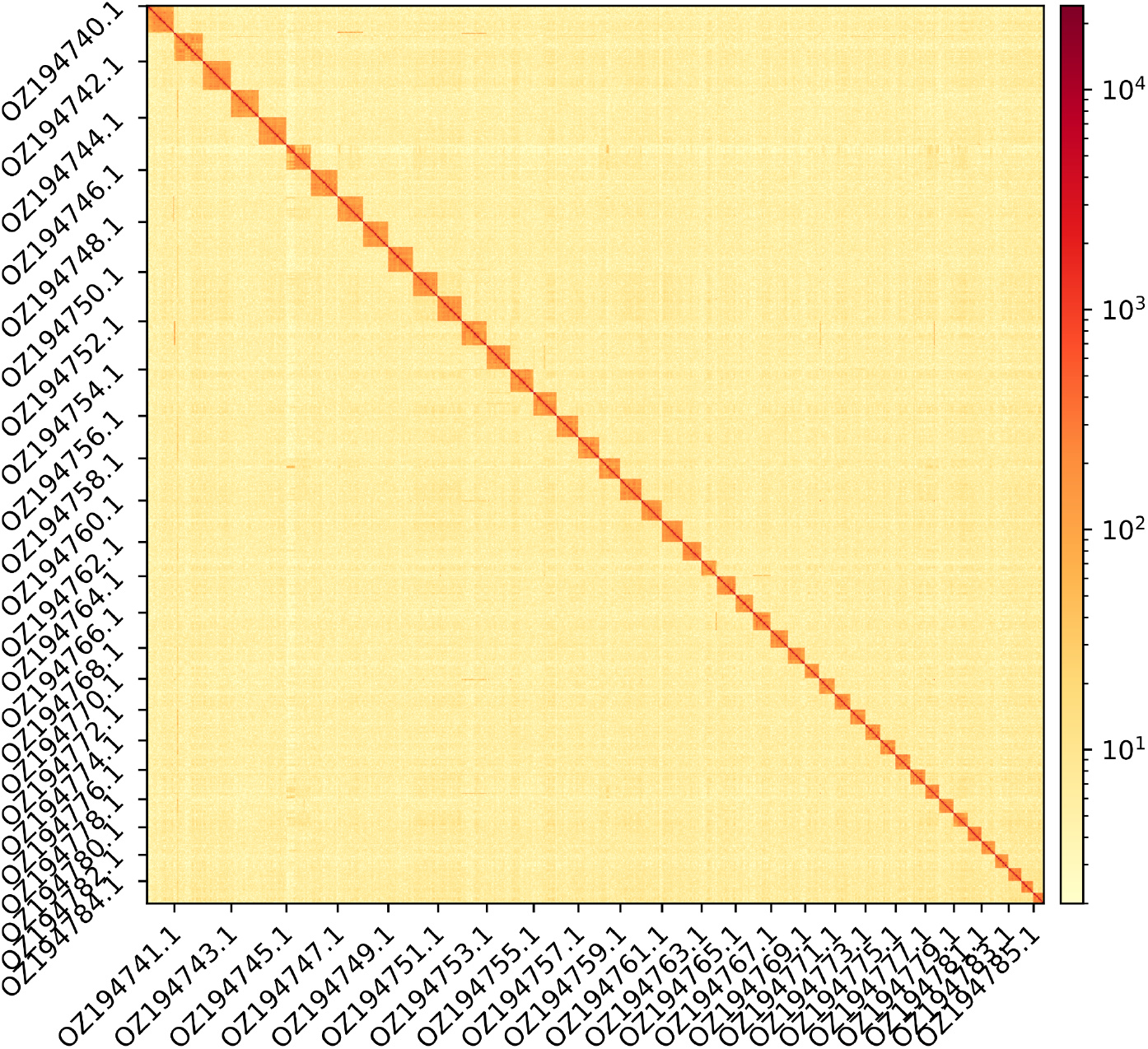
Hi-C contact map showing spatial interactions between regions of the genome. The diagonal corresponds to intra-chromosomal contacts, depicting chromosome boundaries. The frequency of contacts is shown on a logarithmic heatmap scale. Hi-C matrix bins were merged into a 25 kb bin size for plotting. Names of alternate chromosomes have been labelled on x- and y-axes for visualisation purposes.

## Acknowledgements

The specimen was captured during the oceanic research cruise DESAFIO (Ref. PID2020-118118RB-I00). We would like to acknowledge the assembly reviewer, Adama Ndar, from Genoscope. The authors acknowledge the support of the Freiburg Galaxy Team: Saim Momin and Björn Grüning, Bioinformatics, University of Freiburg (Germany), funded by the German Federal Ministry of Education and Research BMBF grant 031 A538A de.NBI-RBC and the Ministry of Science, Research and the Arts Baden-Württemberg (MWK) within the framework of LIBIS/de.NBI Freiburg.

## Conflict of Interest

The authors declare no conflict of interest related to this study. The funding sources had no involvement in the study design, collection, analysis, or interpretation of data; in the writing of the manuscript; or in the decision to submit the article for publication. All authors have participated sufficiently in the work to take public responsibility for the content and agree to the submission of this manuscript.

## Funder Information

This project received funding from Horizon Europe under the Biodiversity, Circular Economy and Environment (REA.B.3); co-funded by the Swiss State Secretariat for Education, Research and Innovation (SERI) under contract numbers 22.00173 and 24.00054; and by the UK Research and Innovation (UKRI) under the Department for Business, Energy and Industrial Strategy’s Horizon Europe Guarantee Scheme.

Funding was provided by the Spanish Ministry of Science, Innovation and Universities (OCTOSET, Ref. RTI2018-097908-B-I00; ECOPHYN, Ref. PID2021-126824NB-C32; MCIU/AEI/FEDER, EU), and the Spanish government through the “Severo Ochoa Center of Excellence” accreditation (CEX2019-000928-S). F.Á.F.-Á. was supported by a Beatriu de Pinós fellowship from Secretaria d ‘Universitats i Recerca del Departament de Recerca i Universitats of the Generalitat de Catalunya (Ref. BP 2021 00035).

## Author Contributions

AB-B collected the samples, FÁF-Á identified the species, FÁF-Á sampled and preserved biological material and provided metadata, MC and AR provided material and information for vouchering and barcoding, AsB provided sampling and metadata support and management, RM, NE, RF, and AsB provided support in sampling, shipping of biological material, metadata collection, and management, LA and MG extracted DNA, prepared libraries, and performed sequencing, FCF, FC and JGG performed genome assembly and curation under the supervision of TSA, LH, and FM performed genome annotation, TB generated the analysis and report. All authors contributed to the writing, review, and editing of this genome note and read and approved the final version.

## Notes

https://www.ebi.ac.uk/ena/browser/view/PRJEB81320

### Competing Interest Statement

The authors have declared no competing interest.

